# A protocol for the reduction of applied torque on parent vessel during elastase-induced aneurysm formation using rabbit animal models

**DOI:** 10.1101/2021.10.08.463749

**Authors:** Sergio A. Pineda-Castillo, Keely A. Laurence, Hannah Homburg, Kar-Ming Fung, Bradley N. Bohnstedt, Chung-Hao Lee

## Abstract

Endovascular therapies for intracranial aneurysms requires animal models for testing the safety and effectiveness prior to translation to the clinic. Rabbits combined with the elastase and right common carotid artery (RCCA) ligation methods is currently a widely used animal model for endovascular device testing. However, the injection of elastase utilizing angiocatheters may potentially exerts adverse torque to the parent vessel and the optimal aneurysm creation period has not been well investigated. In this study, we present a modification to the elastase/RCCA-ligation method by replacing the angio-catheter with a butterfly catheter. Formation of saccular aneurysms was introduced in New Zealand white rabbits (*n* = 6), and were maintained for 2, 4 and 6 weeks. The formed aneurysms exhibited an elongated geometry and were stable during the study period. We found that the modification in the animal surgery procedure provides improved manipulation of the surgical area, prolonged injection of elastase, and effective degradation of the vascular elastic lamina. Compared to the traditional elastase/RCCA-ligation method, the present technique can more effectively reduce unwanted injury to the parent vessel and, therefore, improved stability of the vasculature for testing the efficacy of newly developed endovascular embolization devices.

## 1. Introduction

Subarachnoid hemorrhage (SAH) is a type of stroke that is caused by the rupture of an intracranial aneurysm (ICA), and can lead to death or long-term disability. Currently, the average incidence of aneurysm-provoked SAH is still at 6.1 (95% CI, 4.9-7.5) per 100,000 person-years, in spite of the decreased worldwide prevalence of smoking and high-blood pressure,^[1]^ and the relatively low risk of ICA rupture (0.54 − 1.3% annually).^[2,3]^ However, ICA rupture risk can increase with multiple cofactors, including sex (females being at higher risk than males), Japanese or Finish descent, aneurysm morphological characteristics and location, among others.^[4,5]^ Specifically, geometrical and location factors lead to a wide variety of aneurysm types, which will determine the treatment of choice.

Current ICA treatment methods include microsurgical clipping and endovascular therapies. Micro-surgical clipping utilizes a platinum clip to block blood flow into the ICA, which is highly invasive due to the need of craniotomy. On the other hand, endovascular devices transport occlusive materials into the aneurysm using catheters, which results in reduced in-hospital fatalities, disability and rebleeding, when compared to microsurgical clipping.^[6]^ Therefore, endovascular devices are the preferred method by physicians for aneurysm therapeutics, especially for the elderly and/or patients with aneurysms on the non-middle cerebral artery.^[7]^

Endovascular aneurysm therapy was first introduced with the development of Gugliemi’s detachable coils (GDCs),^[8,9]^ a system that utilizes platinum coils to occlude the aneurysm space and prevent intra-aneruysmal flow. Ever since GDCs introduction, several assisting devices, such as balloons and stents, have been developed, further extending the applicability of the system.^[10]^ However, GDCs face limitations such as mass effects,^[11]^ rebleeding,^[12]^ incomplete occlusion and aneurysm recurrence.^[13]^ This has led to the need for the development of endovascular devices that can overcome the limitations and suboptiomal outcomes of GDCs (e.g., coils with dynamic stiffness,^[14]^ hydrogel-coated coils,^[15,16]^ flow diverters,^[17]^ self-expanding nitinol meshes^[18,19]^).

Despite the wide variety of currently available endovascular devices, aneurysm recurrence is still an emerging challenge for the field of endovascular treatment for ICAs. Therefore, different tools have emerged to assess the degree of occlusion provided by endovascular embolization procedures, such as the Raymond-Roy occlusion classification (RROC). The RROC is a clinical standard for angiographic assessment of the degree of occlusion of an aneurysm: RROC-I represents a completely occluded aneurysm; RROC-II represents a lack of occlusion at the neck of the aneurysm (i.e., neck remnant); and RROC-III represents flow of angiographic dye into the aneurysm dome (**Fig. 1**). **Table 1** lists RROC rates achieved after treatment with different marketable endovascular devices, showing that a maximum of 54.4% patients treated with endovascular methods will have complete occlusion and that the rates of neck and aneurysm remnant are still limited. Specifically, considering that an RROC-III has been demonstrated to pose a substantial risk for aneurysm recurrence and retreatment,^[20]^, it can be inferred that the *state-of-the-art* devices for endovascular embolization cannot prevent aneurysm recurrence. Hence, it is of paramount importance that the newly developed endovascular devices aim to achieve *complete and sustained* occlusion of ICAs (represented by RROC-I), as much as possible.

**Table 1:** Raymond-Roy Occlusion Classification rates after treatment with current marketable endovascular devices.

| Device | RROC Scale |  |  |
| --- | --- | --- | --- |
|  | I | II | III |
| GDCs<sup>[13]</sup> | 25.7% | 32.3% | 42% |
| Hydrogel Coils<sup>[15]</sup> | 54.4% | 19.7% | 25.9% |
| Woven EndoBridge<sup>[18]</sup> | 53.8% | 30.8% | 15.4% |
| Flow Diverters<sup>[24]</sup> | 53.8% | 3.8% | 11.5% |

**Figure 1:**
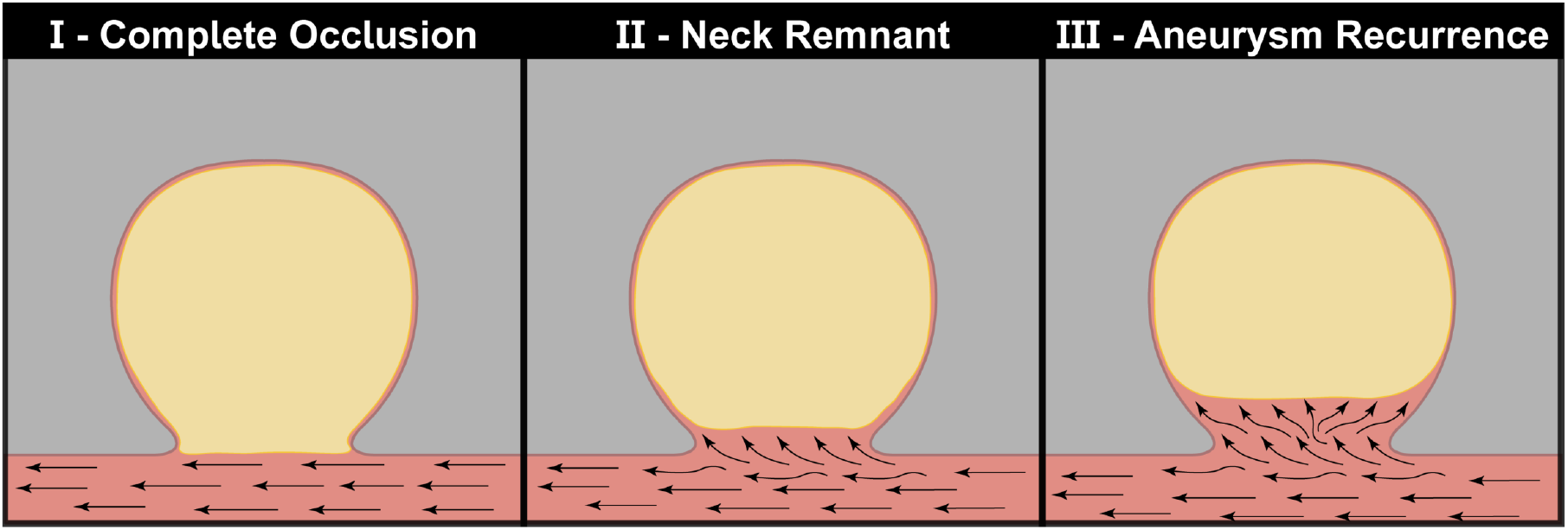
Representation of RROC scale for the assessment of the degree of occlusion of saccular aneurysms.

With this in mind, the development of biomedical devices requires careful and thorough testing before its translation to the clinic. Specifically, biomedical technology requires the use of animal models that serve as platforms to test the *in vivo* safety and efficacy of the device. Thus, modeling of ICAs in animals is of fundamental importance for the development of endovascular devices. Animal models for ICA research have been developed for pathological and biological studies,^[21]^ drug testing^[22]^ and device translation.^[23]^ These include mice, rat, rabbit, dog and pig models, where rabbit and rat carotid artery models are the most frequently used.

ICA formation in these animal models can be performed via different methods, including parent artery ligation, elastase injection, among others. (For a review in ICA animal models, see Strange *et al*.^[25]^) These methods and their combinations are widely accepted for *in vivo* testing of endovascular devices. For example, Killer *et al*.. (2010) used a combination of right common carotid artery ligation (RCCA) and elastase injection to create aneurysms in rabbits for the testing of a hydrogel-coated coil. Also, Herting *et al*. (2020) and Ding *et al*. (2016) used similar methods to test a polymer-coated coil and a self-expanding double-layer nitinol mesh for endovascular embolization, respectively.^[26,27]^

In this study, we aimed at improving the execution of the elastase/RCCA-ligation method for the rabbit aneurysm model. Our findings suggest that this modification will improve the surgical procedure for the formation aneurysms and, ultimately, facilitate a more streamlined procedure for testing ICA embolic devices.

## 2. Methods

### 2.1. Aneurysm Formation

Experimental elastase aneurysms were formed in male New Zealand white rabbits (Charles River Laboratories, NY, USA) (weight: 3.0 − 4.0 kg). Rabbits were sedated with an intramuscular injection of Ketamine (50mg*/*kg) and Xylazine (5mg*/*kg). Sedation was maintained using 1 − 4% isofluorane inhalation. The RCCA was exposed by placing the anesthesized rabbit in supine position and making an incision between the medial edge of the sternocleidomastoid and the trachea. A clip was tightly secured around the RCCA, close to the brachiocephalic trunk. A silk tie was placed around the artery and tied securely at 1.5cm above the aneurysm clip. A 1 : 2 mixture of elastase (BD Millipore) and saline was injected into the RCCA while maintaining the aneurys clip secured. After 20 minutes the aneurysm clip was removed to restore flow. (**Fig. 2a**). Elastase solution enzymatically degraded the healthy elastic lamina (**Fig. 2b**.**1-2**), which debilitated the wall and induced ballooning of the arterial wall (**Fig. 2b**.**3**). The incision was closed and covered with triple antibiotic ointment. The rabbit was then moved to a recovery chamber for recover, and monitored daily for 7 days post-surgery.

**Figure 2:**
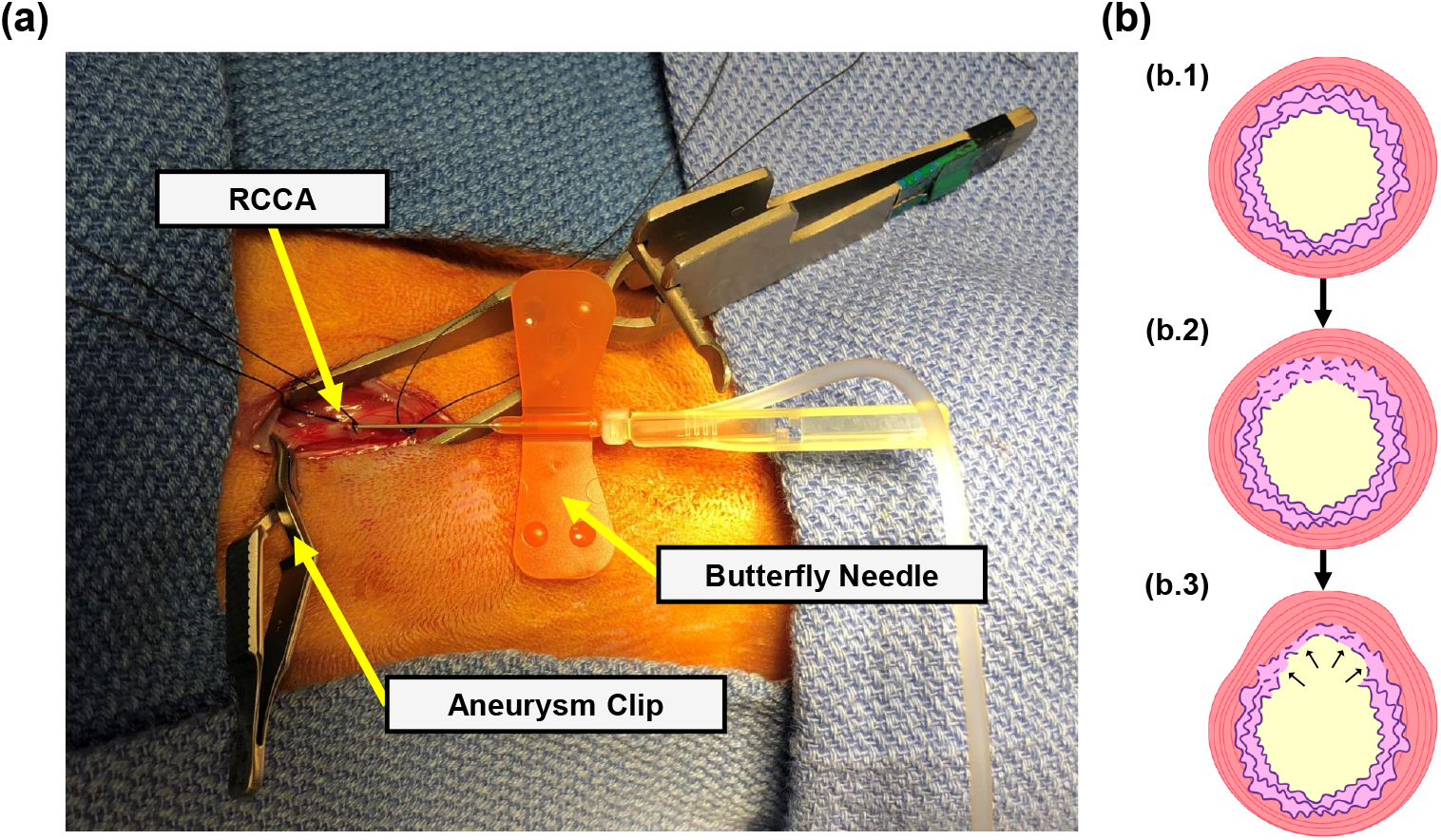
**(a)** Surgical setup for aneurysm formation, showing the use of the butterfly catheter. **(b)** Schematic of aneurysm formation via elastase injection: **(b**.**1)** healthy vascular wall with integral elastic lamina, **(b**.**2)** elastic lamina degradation induced by elastase injection and **(b**.**3)** focal dilation of the arterial wall caused by the breakdown of the elastic lamina.

### 2.2. Assessment of Aneurysm Formation

Rabbits were prepared for angiography at the end of 2, 4, and 6 weeks (*n* = 4, per group) periods using the same anesthetic procedures described previously. The left femoral artery was exposed and an aneurysm clip was placed proximally and the artery was tied about 2cm distal to the clip. A small cut was performed and the catheter and the guide wire were inserted. A C-arm angiography device (GE 9800 Omnipaque contrast) was used to confirm that the catheter reached the ascending aorta. Digitally subtracted angiography was then performed to demonstrate the aneurysm. Imaging was performed to assess aneurysm growth and maintenance. After aneurysm visualization, the catheter was positioned in the left ventricle and Euthasol (1mL*/*4.5kg) were injected. After confirmation of death, the original incision was re-opened to expose the RCCA and resect the aneurysm. The left common carotid artery was also resected as control. Tissues were stored in saline prior to further processing.

Angiographic images were processed to measure aneurysm length (*AL*) and aneurysm width (*AW*), using open-source software ImageJ.^[28]^ Measurments were compared at the three imaging timepoints using a one-way ANOVA test. Differences were deemed significant if *p <* 0.05.

### 2.3. Histology

Resected aneurysms tissues and controls were processed with routine histology methods. Briefly, tissues were fixed in 10% formalin and embedded in paraffin. Tissue sections of 5 − 7 *µ*m of thickness were obtained using a microtome and stained with Verhoeff-elastic-Van Gieson stain. Tissue sections mounted on glass slides for light microscopy assessment. Digital histology images were obtained using a light microscope. Images were taken using a 4X objective and digitally merged using ImageJ (National Institutes of Health, USA).^[28]^

### 2.4. Statistical Analysis

All quantitative measurements are represented as mean ± standard deviation. Animal groups were compared using ANOVA test, where differences were deemed significant when *p <* 0.05. All statistical assessments were performed in SPSS (IBM Corp.).

## 3. Results

### 3.1. Aneurysm Formation

Male New Zealand white rabbits underwent successful aneurysm creation (*n* = 12). Average aneurysm length was 4.01.9 mm (range 2.2 − 8.4 mm) and an average width of 1.50.4 mm (range 1.0−2.3 mm). **Table 2** lists average aneurysm length and width at the investigated timepoints. ANOVA analysis returned no significant difference for length (*p* = 0.983) or width (*p* = 0.401). **Fig. 3** shows representative angiographic assessment of aneurysm formation at the RCCA.

**Table 2:** Raymond-Roy Occlusion Classification rates after treatment with current marketable endovascular devices.

| Timepoint | AL (mm) | AW (mm) |
| --- | --- | --- |
| 2 weeks | $4.2 \pm 2.8$ | $1.3 \pm 0.3$ |
| 4 weeks | $3.9 \pm 1.0$ | $1.4 \pm 0.2$ |
| 6 weeks | $3.9 \pm 1.8$ | $1.7 \pm 0.4$ |

**Figure 3:**
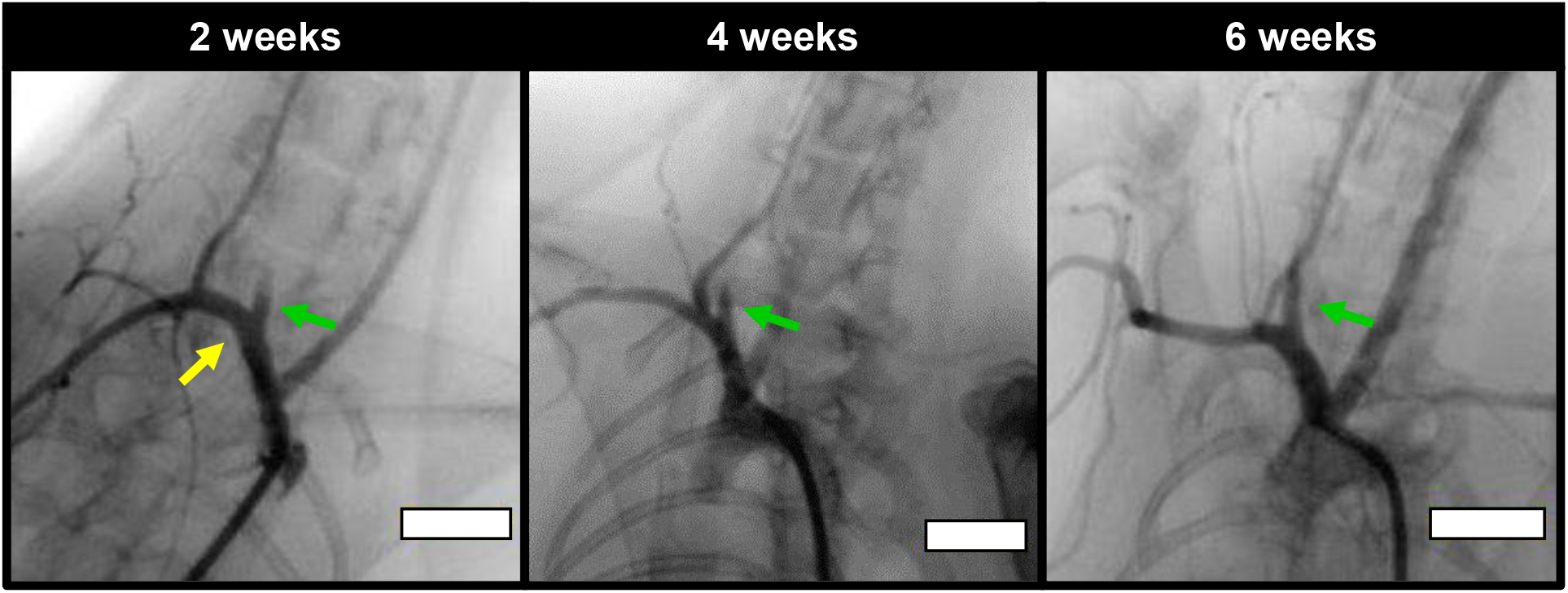
Representative angiograms of the created aneurysms. The green arrows point towards the created aneurysms and the yellow arrow indicates the RCCA. Scale bars = 10 mm)

### 3.2. Histological Assessment

Histological sections were succesfully obtained from the resected aneurysms. Tissue sections exhibited the classic structure for the arterial wall of the aneurysm. Specifically, Verhoeff-Van Gieson stain allowed the observation of the elastic fibers in the vascular wall, which were observed as the darker purple tone in **Fig. 4**. Our results indicate a generalized deficiency of elastic fibers in the aneurysm wall, where we even observed regions sections with complete absence of these elastic structures. Additionally, we observed that the aneurysms of the 6-week group exhibited an evident lack of structure in the elastic fibers (**Fig. 4c**), while the other two time points still exhibited some fragments elastic lamina (**Fig. 4a**,**b**).

**Figure 4:**
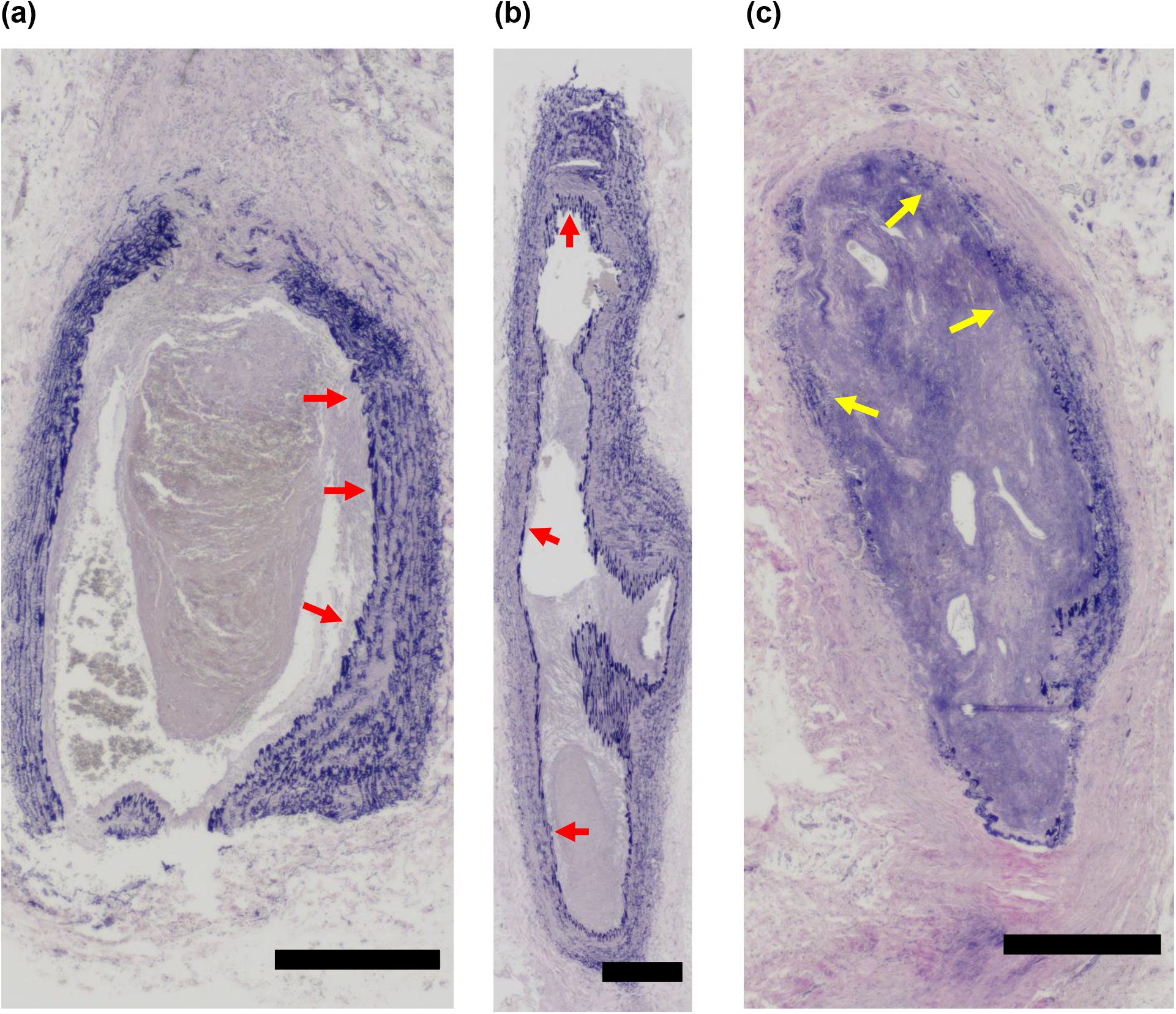
Histology slides of the created aneurysms depicting elastin degradation at **(a)** 2 weeks, **(b)** 4 weeks, and **(c)** 6 weeks. Red arrows indicate the portions of the non-degraded elastic lamina. Yellow arrows indicate the degradation of the elastic fibers. Scale bars = 0.5 mm.

## 4. Discussion

### 4.1. Our approach

Technology for endovascular embolization of intracranial aneurysms has advanced significantly over the last thirty years since the development of GDCs. Further innovation in the field requires the development of pre-clinical models that allow testing of safety and efficacy of endovascular devices. To this end, animal models for saccular aneurysms have been developed and improved over the past six decades.^[29–31]^ These models have been able to accurately mimic the anatomy and hemodynamics of human aneurysms using different species, including mice, rats, rabbits, among several others. Specifically, the rabbit model has proved to be superior for the testing of endovascular devices, due to the docility of the animal, low maintenance cost, wide availability and appropriate size for the formation of aneurysms with similar geometry to the human.^[25]^

Among the different aneurysm formation techniques in the rabbit model, elastase injection is described as a fast alternative for the formation of aneurysms prone to rupture.^[25]^ In this procedure, successful elastase injection requires proximal and distal ligation of the common carotid artery (CCA), which is consistently described to included the dissection of the parent artery and the use of a silk tie for distal ligation of the CCA. Currently, two techniques for proximal ligation of the vessel are extensively applied: (i) the use of a Fogarty balloon^[32]^, or (ii) use of a temporary aneurysm clip.^[33]^ Based on our experience, we agreed with Hoh *et al*. about the potential damage to the CCA when using the Fogarty balloon. Therefore, we have adapted the method as described by Hoh *et al*. and aimed at its improvement.

Hoh *et al*. used of an angiocatheter to inject elastase between the proximal and distal ligation sites and it was maintained in the intraluminal space for the 20-minute elastase injection time. We initially performed our aneurysm formation with the angiocatheter but we noticed that it had the potential to cause torque on the vessel. In addition, it was not only more difficult to insert the catheter into the CCA initially, but it placed unwanted pressure on the sides of the arterial wall. These complications during elastase injection might lead to unwanted vessel lesion and and remodeling processes on the artery, ultimately altering the native condition of the healthy artery wall.^[34]^ As a result, we have substituted the use of the angiocatheter for a butterfly catheter to reduce the potential application of torque. This modification of the protocol allowed the surgical team to to step back from the table and inject elastase from the opposite end of the catheter without risk of vessel damage. As a result, we have been able to induce the formation of stable saccular aneurysms via elastic lamina degradation, as demonstrated in **Fig. 3** and **Fig. 4**. We believe that this modification to the aneurysm formation protocol can provide significant improvement to the ease-of-use of this animal model.

### 4.2. Applications for Endovascular Device Testing

Formation of saccular aneurysms for the testing of endovascular devices requires the maintenance of native properties in the parent artery, as off-target damages during elastase injection might lead to unwanted vessel rupture or reduce effectiveness of the model, in addition to animal fatalities. In addition, damages to the vessel wall might also increase alterations in the vessel flow dynamics, limiting the accuracy of aneurysm computational flow dynamics research, which is paramount importance for the development of endovascular devices.^[35]^

Our findings in this modified protocol will be applied to the development of a novel coil-free endovascular device. We have developed a shape memory polymer (SMP) for endovascular embolization of patient-specific aneurysm geometries,^[36–39]^ and we believe that this elastase-induced aneurysm formation protocol can aid in the translation of our material to the pre-clinical setting. We plan to test our material with a statistically powerful sample size, to determine the pre-clinical safety and effectiveness of our SMP device (measured using RROC scale, **Fig. 1**). We also plan on studying the microstructural and mechanical properties of the aneurysms formed in this animal model, and compare them to our recent findings in the properties of a human aneurysm specimen.^[40]^ Lastly, this animal model can also be used for training of surgical teams in the deployment of current marketable endovascular devices.^[41]^

### 4.3. Study Limitations

The key advantage of this technique over similar techniques is the significant reduction in the torque on the artery. While this is not necessarily stated as an issue in the literature, we experienced vessel torsion while using an angiocatheter and injecting the elastase solution. To minimize the risk of injury, it was imperative that we substitute the angiocatheter, which was done by using a butterfly catheter. We also found that, while the balloon catheter method was effective, this step required more time than was necessary to achieve the same result. While we did not encounter any short term complications from our procedure, or any post-operative complications with any of the animals, it would be advantageous to replicate this protocol with more samples and/or a longer study period.

## 5. Conclusion

In this study, we present a modification to traditional elastase-induced aneurysm models. We have substituted the use of an angiocatheter for elastase injection for a butterfly catheter, which reduced the applied torque on the artery, reducing the potential for parent vessel injury. We believe that this modification to the traditional elastase/RCCA-ligation method can further improve aneurysm modeling in rabbit animal models. This method improvement has important implications for the *in vivo* testing of endovascular devices and training for ICA embolization procedures.

## Acknowledgments

We would like to acknowledge the supported granted by the Oklahoma Center for the Advancement of Science and Technology (OCAST) Health Research Program (HR18-002), the Oklahoma Shared Clinical and Translational Resources (OSCTR) Pilot Projects Program (NIGMSU54GM104938), and the institutional funding from the VPRP Office and the Institute for Biomedical Engineering, Science and Technology (IBEST) at the University of Oklahoma.

## Conflict of Interest

*The authors declare no conflict of interest in the presented work*..

## Notes

### Competing Interest Statement

The authors have declared no competing interest.

